# Draft genome sequences for *Enterococcus mundtii* isolates from soil in Minnesota

**DOI:** 10.1101/2025.11.06.687061

**Authors:** Erna Karic, Amanda L. Haeberle, Julia L. E. Willett

## Abstract

*Enterococcus mundtii* is a Gram-positive bacterium found in the environment and mammalian gastrointestinal tracts. It is also an infrequent cause of infections in humans. Here, we describe isolation, sequencing, and genomic analysis of four isolates of *E. mundtii* obtained from soil. These strains will be valuable tools for future work on understuded species of *Enterococcus*.

## Results

*Enterococcus* species are frequently found as gastrointestinal tract commensals in mammals, birds, and insects, and these bacteria are prevalent in the environment (1-4). The best-studied enterococcal species, *E. faecalis* and *E. faecium*, cause infections in humans and can be highly resistant to antimicrobials (5). Other enterococci are relatively understudied yet display interesting phenotypes such as pigmentation and motility (6, 7). This study presents the draft genome sequences of 4 isolates of *Enterococcus mundtii*, a pigmented non-motile bacterium (8, 9). Soil samples were collected in collaboration with the MICB 4215 course at the University of Minnesota-Twin Cities during October 2023. Approximately 0.5 grams of soil were resuspended in 0.5 mL sterile water, and 0.1 mL was plated on both tryptic soy agar (TSA) and bile esculin azide (BEA) agar, a selective medium for enterococci. Plates were incubated for 24 h at 37 °C. Four presumptive *Enterococcus* isolates (dark colonies on BEA plates) were obtained from a soil sample taken after a rain storm from beneath a large mushroom near the University of Minnesota. These four colonies, designated 4215-23-A1 through D1, were chosen for further analysis due to their yellow pigmentation on TSA plates, which differs from other enterococci such as *Enterococcus faecalis*. All putative enterococci were restreaked, grown in tryptic soy broth (TSB), and stored at −80 °C in TSB supplemented with 25% glycerol.

Samples from freezer stocks were grown static in TSB for 24 h at 37°C. DNA was isolated using a Qiagen DNeasy Blood and Tissue kit with a lysozyme pre-treatment as previously described for *E. faecalis* (10). Sample purity and concentration were determined using a NanoDrop and Qubit fluorometer, respectively. Library preparation and whole-genome sequencing were performed at SeqCoast. Libraries were prepared with the Illumina DNA Prep tagmentation kit and IDT For Illumina Unique Dual Indexes. An Illumina NextSeq2000 (300 cycle flow cell, 2×150bp paired-end reads) was used for sequencing. DRAGEN (v3.10.12) was used for demultiplexing and trimming. These reads were assembled in Unicycler v0.4.8 (11) and annotated using RASTtk (12) in the Bacterial and Viral Bioinformatics Resource Center (BV-BRC, https://www.bv-brc.org/). Samples were identified as *E. mundtii* using the BV-BRC Taxonomic Classification Service, and CheckM was used to assess genome quality and completeness. This classification was consistent with colony phenotypes, specifically production of yellow pigment.

*E. mundtii* 4215-23-A1 has a genome length of 3,122,934 bp, which was assembled into 67 contigs (N50 length = 237,970 bp) with 99x coverage and 3,037 protein-coding sequences. *E. mundtii* 4215-23-B1 has a genome length of 3,040,745 bp assembled into 53 contigs (N50 length = 238,061 bp) with 102x coverage and 2,939 protein-coding sequences. *E. mundtii* 4215-23-C1 is 3,051,831 bp in length and was assembled into 59 contigs (N50 length = 238097 bp) with 135x genome coverage and 2,953 protein-coding sequences. *E. mundtii* 4215-23-D1 has a genome length of 3,052,711 bp, with 54 contigs (N50 length = 279,319 bp), 127x genome coverage, and 2,954 protein-coding sequences. All strains had 49 tRNAs and 2 16S rRNA genes. The GC content ranged from 38.23-38.35%, similar to other *E. mundtii* strains. Cladogram analysis (**Figure 1**) revealed that our isolates were closely related to *E. mundtii* strain 4A2-18, a soil isolate from South Africa (GenBank assembly GCF_015230405.1).

**Figure 1.**
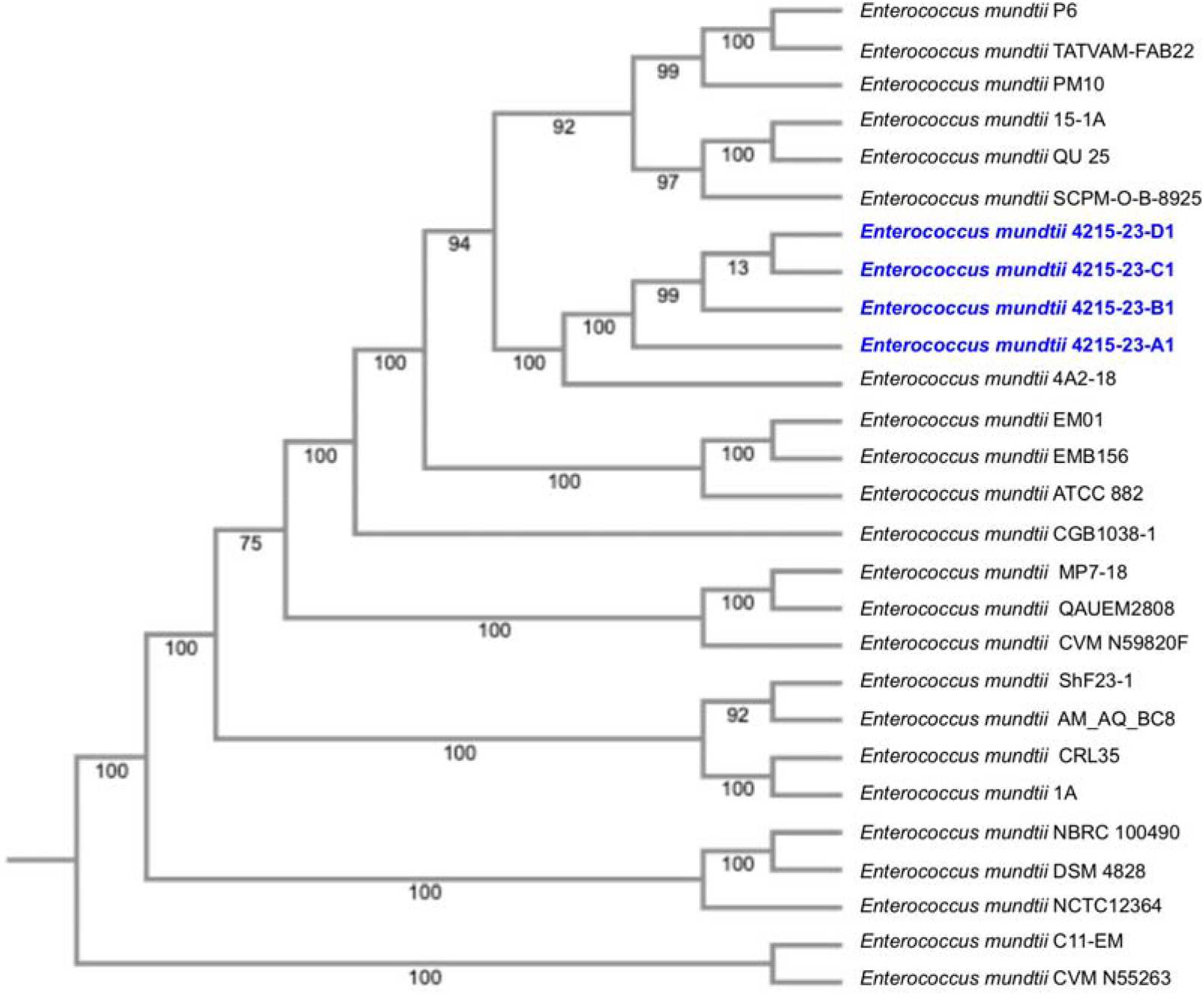
Comparison of *E. mundtii* isolates described in this study to existing isolates. The cladogram was generated using RAxML v8.2.12 with 100 rounds of fast bootstrapping. Strains from this study are indicated with bold, blue text.

## Data availability

Raw reads from these isolates were deposited with BioProject number PRJNA1345430.

## Acknowledgements

We thank the students in the University of Minnesota College of Biological Sciences course MICB 4215 during fall 2023 for their enthusiasm. ALH was supported in part by National Institutes of Health training grant T32GM140936.

## References

1. Lebreton F, Manson AL, Saavedra JT, Straub TJ, Earl AM, Gilmore MS. Tracing the Enterococci from Paleozoic Origins to the Hospital. Cell. 2017;169(5):849-61.e13.

2. Schwartzman JA, Lebreton F, Salamzade R, Shea T, Martin MJ, Schaufler K, et al. Global diversity of enterococci and description of 18 previously unknown species. Proc Natl Acad Sci U S A. 2024;121(10):e2310852121.

3. Byappanahalli MN, Nevers MB, Korajkic A, Staley ZR, Harwood VJ. Enterococci in the environment. Microbiol Mol Biol Rev. 2012;76(4):685–706.

4. Goh HMS, Yong MHA, Chong KKL, Kline KA. Model systems for the study of Enterococcal colonization and infection. Virulence. 2017;8(8):1525–62.

5. Willett JLE, Dunny GM. Insights into ecology, pathogenesis, and biofilm formation of Enterococcus faecalis from functional genomics. Microbiol Mol Biol Rev. 2025;89(1):e0008123.

6. Palmer KL, Godfrey P, Griggs A, Kos VN, Zucker J, Desjardins C, et al. Comparative genomics of enterococci: variation in Enterococcus faecalis, clade structure in E. faecium, and defining characteristics of E. gallinarum and E. casseliflavus. MBio. 2012;3(1):e00318–11.

7. Aarestrup FM, Butaye P, Witte W. Nonhuman Reservoirs of Enterococci In: Michael S. Gilmore DBC, Patrice Courvalin, Gary M. Dunny, Barbara E. Murray, Louis B. Rice, editor.: ASM Press; 2002.

8. Ran Q, Badgley BD, Dillon N, Dunny GM, Sadowsky MJ. Occurrence, genetic diversity, and persistence of enterococci in a Lake Superior watershed. Appl Environ Microbiol. 2013;79(9):3067–75.

9. Repizo GD, Espariz M, Blancato VS, Suárez CA, Esteban L, Magni C. Genomic comparative analysis of the environmental Enterococcus mundtii against enterococcal representative species. BMC Genomics. 2014;15(1):489.

10. Willett JLE, Dale JL, Kwiatkowski LM, Powers JL, Korir ML, Kohli R, et al. Comparative Biofilm Assays Using Enterococcus faecalis OG1RF Identify New Determinants of Biofilm Formation. mBio. 2021;12(3):e0101121.

11. Wick RR, Judd LM, Gorrie CL, Holt KE. Unicycler: Resolving bacterial genome assemblies from short and long sequencing reads. PLoS Comput Biol. 2017;13(6):e1005595.

12. Brettin T, Davis JJ, Disz T, Edwards RA, Gerdes S, Olsen GJ, et al. RASTtk: a modular and extensible implementation of the RAST algorithm for building custom annotation pipelines and annotating batches of genomes. Sci Rep. 2015;5:8365.

